# Differential pathogenesis of Usutu virus isolates in mice

**DOI:** 10.1101/2020.08.31.275149

**Authors:** Sarah C. Kuchinsky, Seth A. Hawks, Eric Mossel, Sheryl Coutermarsh-Ott, Nisha K. Duggal

## Abstract

Usutu virus (USUV; *Flavivirus*), a close phylogenetic and ecological relative of West Nile virus, is a zoonotic virus that can cause neuroinvasive disease in humans. USUV is maintained in an enzootic cycle between *Culex* mosquitoes and birds. Since the first isolation in 1959 in South Africa, USUV has spread throughout Africa and Europe. Reported human cases have increased over the last few decades, primarily in Europe, with symptoms ranging from mild febrile illness to severe neurological effects. In this study, we investigated whether USUV has become more pathogenic during emergence in Europe. Interferon α/β receptor knockout (*Ifnar1*^-/-^) mice were inoculated with recent USUV isolates from Africa and Europe, as well as the historic 1959 South African strain. The three tested African strains and one European strain from Spain caused 100% mortality in inoculated mice, with similar survival times and histopathology in tissues. Unexpectedly, a European strain from the Netherlands caused only 12% mortality and significantly less histopathology in tissues from mice compared to mice inoculated with the other strains. Viremia was highest in mice inoculated with the recent African strains and lowest in mice inoculated with the Netherlands strain. Based on phylogenetics, the USUV isolates from Spain and the Netherlands were derived from separate introductions into Europe, suggesting that disease outcomes may differ for USUV strains circulating in Europe. These results also suggest that while more human USUV disease cases have been reported in Europe recently, circulating African USUV strains are still a potential major health concern.

**Author Summary:** Usutu virus (USUV) is an emerging mosquito-borne virus that causes severe neuroinvasive disease in humans. USUV was first detected in Africa in 1959, and cases of human disease have increased in recent years. Most USUV disease cases are now reported in Europe, where the virus currently circulating. One possibility for the increase in case numbers is that USUV strains have become more pathogenic over time during its spread from Africa into Europe. We compared the pathogenesis of five USUV isolates from Africa and Europe in a mouse model. Three isolates from Africa and one isolate from Europe caused 100% mortality in mice. Surprisingly, one isolate from Netherlands caused only 12% motality. Significantly less histopathology was observed in tissues from mice inoculated with the Netherlands strain compared to mice inoculated with the other four strains. Our results suggest that, even though more human USUV disease cases have been reported in Europe recently, African USUV strains are also highly pathogenic.

## Introduction

Usutu virus (USUV) is an emerging zoonotic virus that can cause neuroinvasive disease in humans [1]. As a member of the *Flaviviridae* family, *Flavivirus* genus, USUV is closely related to other pathogens of global human health concern, including West Nile virus (WNV), dengue virus (DENV), yellow fever virus (YFV), and Zika virus (ZIKV) [2]. USUV is typically maintained in an enzootic cycle between ornithophilic mosquito species, including *Culex* spp., and various avian species [3-5]. Incidental or “spillover” infections in mammals, including bats [6], red deer [7], horses, dogs [8], rodents [9], and humans, have been reported [2].

Since the first isolation of Usutu virus (USUV) in South Africa in 1959 from a *Culex neavei* mosquito [10], USUV has spread throughout Africa and Europe [11, 12]. The first emergence of USUV into Europe was observed in Austria in 2001, upon the discovery of the mass mortality of approximately 50,000 Eurasian blackbirds (*Turdus merula*) [13, 14]. USUV outbreaks in Europe have increased over the last two decades, as evidenced by the occurrence of epizootic events in numerous avian species in countries throughout central Europe and surrounding the Mediterranean basin [12]. Phylogenetic analyses estimate that four intercontinental viral migration events of USUV have occurred in the past 60 years [11].

As recently as 2018, there have been 49 documented human cases of acute USUV infection, 47 of which have been reported in Europe [2]. Thirteen cases in Italy [15-18], 6 in Croatia [19, 20], and 1 in France [21] have exhibited severe clinical disease symptoms including: meningoencephalitis, encephalitis, neuroinvasive disease, and idiopathic facial paralysis. Two human clinical cases of USUV infection were reported in Africa, first in 1981 in the Central Africa Republic, and then in 2004 in Burkina Faso [22]; both exhibited mild clinical symptoms, such as febrile illness [2]. USUV antibodies have been detected in healthy blood donors in Serbia, Italy, and Germany [23-25], indicating exposure to USUV. In total, USUV antibodies have been reported in 98 individuals in Europe [2].

Several murine models have been developed to investigate the pathogenesis of USUV. Wild type mice have shown limited development of USUV pathogenesis, unless they are less than one week old. Studies have shown high mortality in suckling mice inoculated with either the prototypic SAAR 1776 strain (South Africa 1959) or a European strain, USUV 939/01 (Vienna 2001) [26, 27]. Suckling mice inoculated with USUV 939/01 exhibited clinical signs including disorientation, paralysis, and coma beginning 6 days post inoculation. Evidence of viral replication in neurons and neuronal apoptosis has been observed histologically in one-week-old mice [27] and indicates the neurovirulence of USUV in young mice.

Studies have also employed immunodeficient murine models to evaluate USUV pathogenesis using a different European strain, USUV V18 (Germany 2011) or the prototypic South Africa 1959 strain. Adult male AG129 mice, deficient in the interferon α/β- and γ-receptors, exhibited clinical signs of disease including rapid weight loss, as well as high mortality rates, when challenged with USUV strain V18 [28]. Martín-Acebes and others reported similar results using adult interferon α/β receptor knockout (*Ifnar1*^*-/-*^) mice, as well as high viral RNA loads in the brain tissues of mice challenged with USUV BIOTEC (derived from the South Africa 1959 strain) [29]. These studies indicate that mice deficient in interferon receptors may serve as appropriate model to study USUV pathogenesis and also suggest the importance of the host interferon response in modulating USUV disease progression.

The previously described murine studies assessed USUV pathogenesis using a single virus strain, either the prototypic South African 1959 strain or a European strain. This study sought to directly compare pathogenesis in an *Ifnar1*^*-/-*^ mouse model elicited by the prototypic South African 1959 strain and 4 recent, low-passage African and European USUV isolates strains: Uganda 2010, Senegal 2003, Spain 2009 and Netherlands 2016, respectively. Given the fewer number of human USUV disease cases reported in Africa, we predicted that Uganda 2010, Senegal 2003, and South Africa 1959 would be less pathogenic than Spain 2009 and Netherlands 2016 in mammals. We found striking differences in mortality and morbidity between USUV isolates in inoculated mice. Furthermore, viral serum titers differed significantly between isolates. Results from this study suggest that USUV pathogenesis is different between strains, with the recent African strains, Uganda 2010 and Senegal 2003, generating higher viremia and pathogenesis in mice than the European strains, Spain 2009 and Netherlands 2016.

## Results

### Differences in morbidity and mortality in mice inoculated with contemporary USUV isolates

Five isolates of USUV were selected for our studies, including the prototypic strain South Africa 1959, as well as four more recent isolates from Africa and Europe: Senegal 2003, Uganda 2010, Spain 2009, and Netherlands 2016 (Table 1). First, we evaluated the growth kinetics of the USUV isolates *in vitro*. All 5 strains replicated similarly in Vero cells (Fig. 1). All strains reached peak viral titer on day post-inoculation (dpi) 2 at approximately 8 log10 PFU/mL.

**Table 1.**

| USUV strain | Country of origin | Year of collection | Species of isolation | Passage number | Citation | Accession number |
| --- | --- | --- | --- | --- | --- | --- |
| SAAR1776 | South Africa | 1959 | *Culex neavei* | p7SM1V1 | [10] | MN813492 |
| DakPM173701 | Senegal | 2003 | *Culex neavei* | pV2+<sup>1</sup> | [22] | MN813488 |
| HU10279-09 | Spain | 2009 | *Culex perixguus* | pV2+<sup>1</sup> | [41] | MN813489 |
| UG09615 | Uganda | 2010 | *Culex* sp. | pV3 | [43] | MN813491 |
| TMNetherlands | Netherlands | 2016 | *Turdus merula* | pV5 | [42] | MN813490 |

<sup>1</sup> unknown complete passage history; SM indicates suckling mouse; V indicates Vero cells

**Figure 1.**
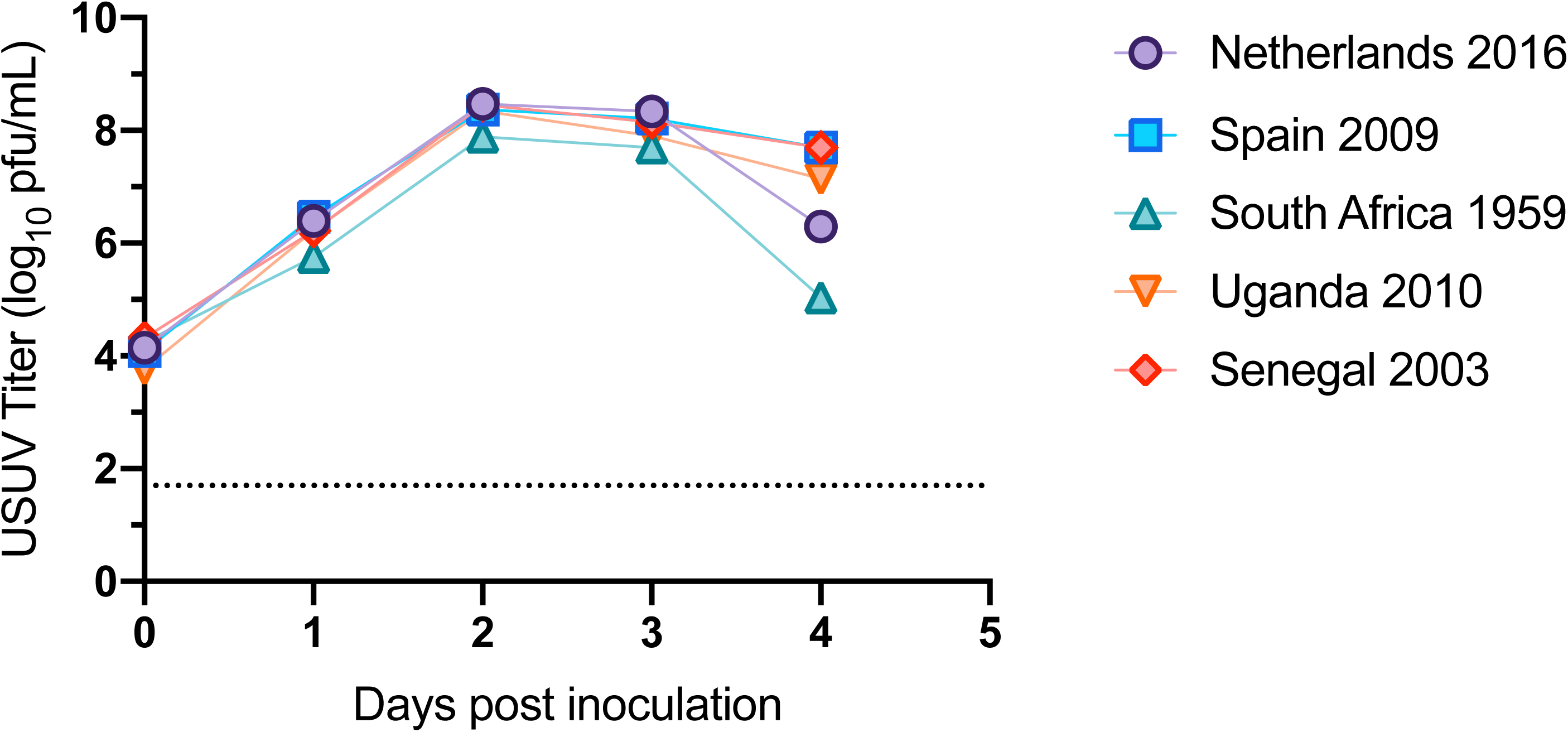
Growth kinetics of USUV isolates in Vero cells. Circles and error bars represent means and standard deviations, respectively, of triplicate inoculated cultures. Dashed line represents limit of detection. The growth curve was performed three times, with one representative experiment shown.

Next, in order to study the virulence of USUV strains, *Ifnar1*^-/-^ mice were subcutaneously inoculated with the five USUV isolates. Survival differed significantly between mice inoculated with the Netherlands 2016 isolate (88%) compared to mice inoculated with the Spain 2009, South Africa 1959, Uganda 2010, and Senegal 2003 isolates (0%) (Fig. 2A, p<0.0001). Mice inoculated with the latter four isolates lost a significant amount of weight compared to mice inoculated with Netherlands 2016 on dpi 5 (Fig. 2B, p<0.0001). Mice that experienced weight loss also exhibited signs such as lethargy and decreased feeding and required euthanasia by dpi 6. Mice inoculated with the Netherlands 2016 isolate did not exhibit extreme weight loss, losing an average of 5% starting weight; by dpi 6, weight gain was observed.

**Figure 2.**
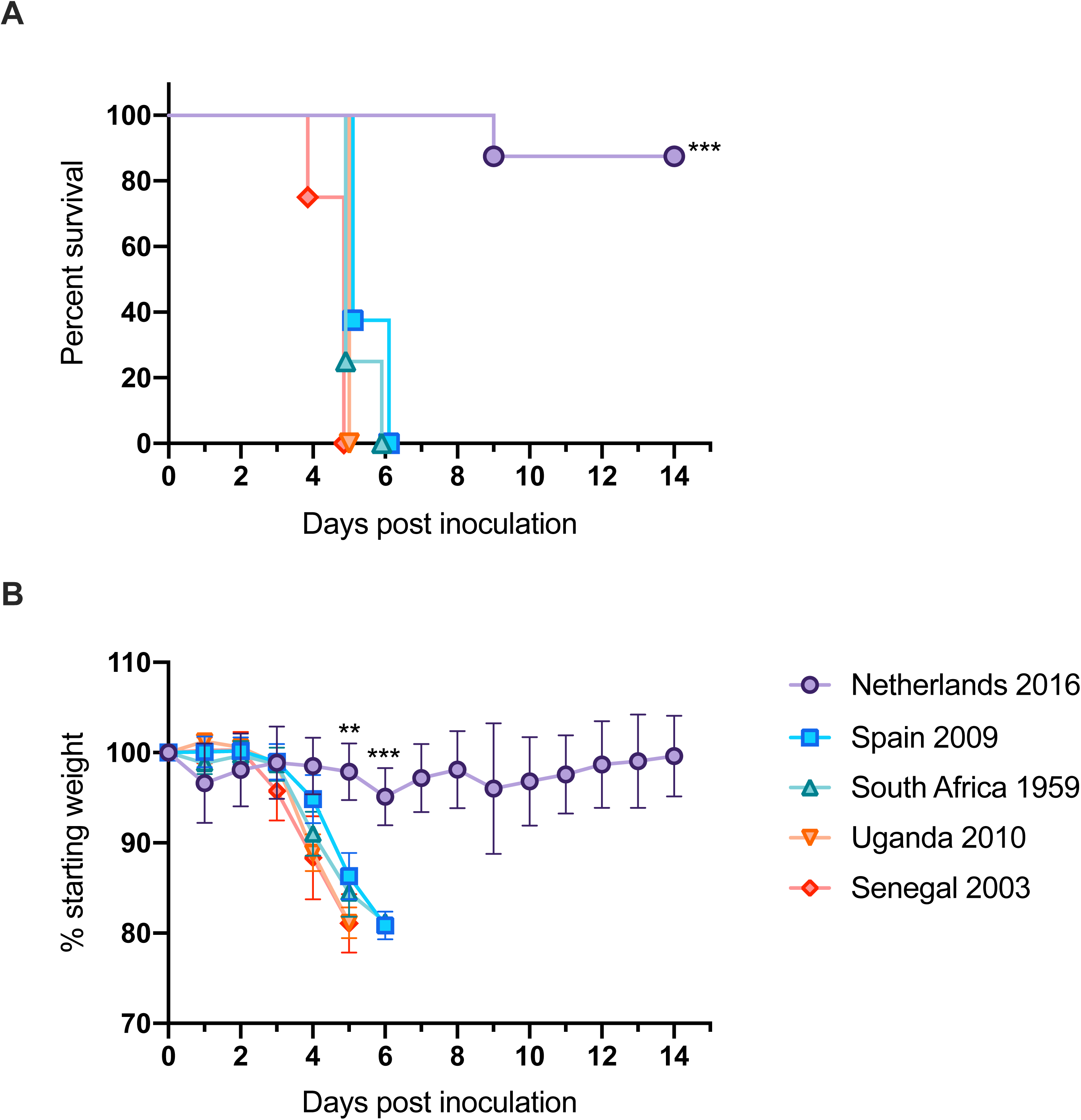
Morbidity and mortality of mice inoculated with USUV isolates. (A) Percent survival of mice post-inoculation. (B) Average percentage of initial weight post-inoculation. ** p<0.01, *** p<0.001.

### Decreased viral dissemination and histopathology in mice inoculated with the Netherlands USUV strain

Mice inoculated with Netherlands 2016 showed significantly lower viremia compared to mice inoculated with the other 4 USUV isolates on all days tested (Fig. 3A, p<0.001). Three distinct viremia profiles were observed: mice inoculated with Netherlands 2016 had the lowest peak titers of 3 log10 USUV pfu/ml; mice inoculated with Spain 2009 and South Africa 1959 had moderate peak titers of 5 log10 USUV pfu/ml of serum; and mice inoculated with Uganda 2010 and Senegal 2003 had the highest peak titers of 7 log10 USUV pfu/ml of serum. Peak titers across all strains were observed on dpi 5.

**Figure 3.**
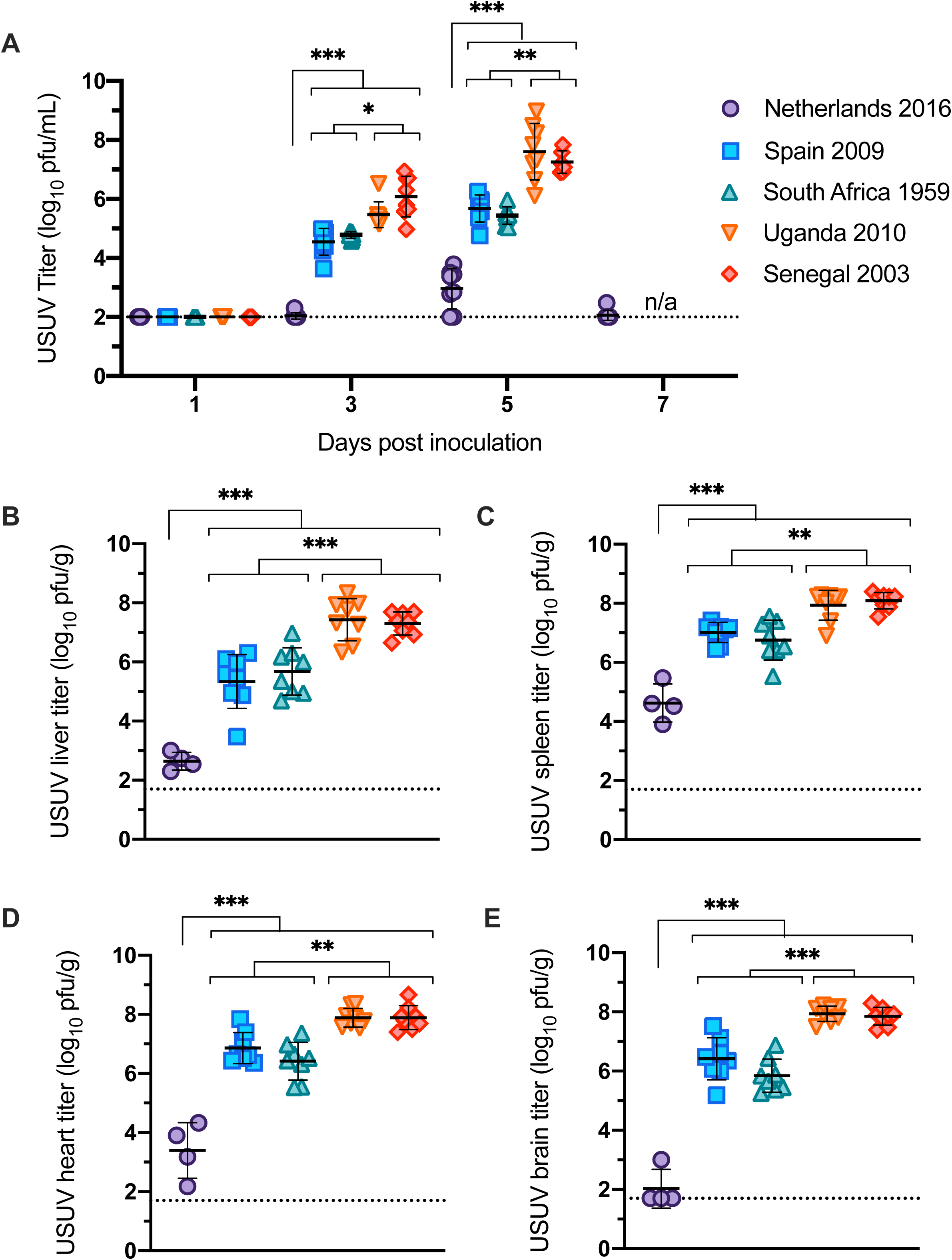
Viral titer of USUV isolates in serum and tissues. (A) Viremia of USUV isolates in mice. Titers are reported as log10 PFU per mL of serum. (B-E) Viral titers of USUV isolates in mouse tissues collected on dpi 5-6. Titers represented as log10 PFU per gram of: (B) liver; (C) spleen; (D) heart; and (E) brain. Lines represent mean; error bars represent standard deviation. Dashed line represents the limit of detection. * p<0.05, ** p<0.01, *** p<0.001.

Significant differences in viral titers were also observed in heart, brain, liver, and spleen tissues (Fig. 3B-E). For all tissues tested, mice inoculated with Netherlands had the lowest viral titer (p<0.001); mice inoculated with isolates from Uganda and Senegal had the highest titer; and mice inoculated with isolates from Spain and South Africa had tissue titers that fell between the other two groups (p<0.01).

To evaluate the pathogenic effects of the USUV strains, sections of brain, spleen, liver, and heart were evaluated microscopically. The most striking changes were identified in the spleen. Spleens of all animals were semi-quantitatively evaluated for inflammation and evidence of cell death in both the white pulp (lymphoid follicles) and red pulp (intervening splenic cords and sinuses). These scores were then summed for a total pathology score. Animals infected with the Spain 2009, Senegal 2003, Uganda 2010, and South Africa 1959 strains of Usutu virus were similar in score and characterized by large amounts of cell death and inflammation in both the red and white pulp (Fig. 4A). Inflammatory cells were predominantly macrophages but often aggregates of neutrophils were scattered throughout as well (Fig. 4C). Spleens from PBS control animals had no significant inflammatory lesions or significant evidence of cell death. Spleens from animals infected with the Uganda 2010 isolate exhibited significant infiltration by macrophages and neutrophils (pictured) as well as prominent cell death (not pictured) within both the red and white pulp. Spleens from animals infected with the Netherlands 2016 isolate looked similarly to the PBS control group, with only minimal evidence of predominantly histiocytic inflammation in the red and white pulp and minimal cell death (Fig. 4C). Additionally, these animals had scores that did not statistically differ from those in the PBS control group.

**Figure 4.**
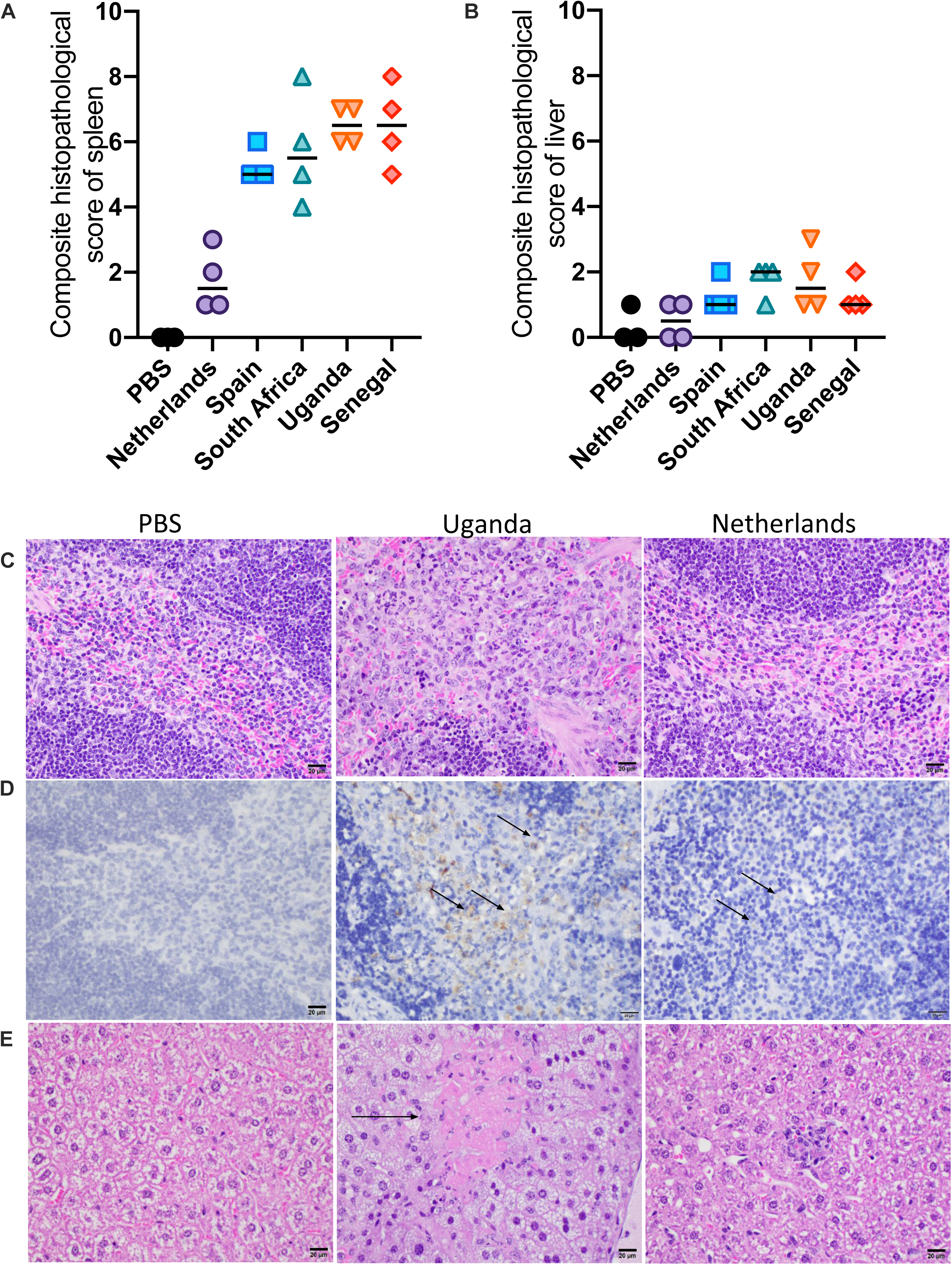
Histopathology of tissues. Samples of spleen, liver, heart, and brain were evaluated microscopically. (A) Composite histopathology scores of spleen tissue; lines represent median. (B) Composite histopathology scores of liver tissue; lines represent median. (C) Spleens from control animals and those infected with the Netherlands strain had no significant inflammatory lesions or significant evidence of cell death. Spleens from animals infected with Uganda, Spain, Senegal, and South Africa strains of Usutu virus exhibited readily observable inflammation and cell death within both the red and white pulp. (D) Immunohistochemistry for Usutu virus antigen revealed high viral loads in macrophages within the spleen of Uganda strain infected animals. Rare antigen was present in macrophages in spleens from those animals infected with the Netherland strain. (E) Livers taken from control animals were characterized by no evidence of inflammation or cell death. Prominent cell death (arrow) and variable inflammation wad identified in the liver tissues of animals infected with Uganda, Spain, Senegal, and South Africa strains of Usutu virus. Mild inflammation was identified in the liver tissues of animals infected with the Netherlands strain. (F) Brains taken from both control animals as well as virus-infected animals showed no microscopic evidence of inflammation or cell death in sections of cerebral cortex.

Sections from liver were also affected in these animals. Livers of all animals were semi-quantitatively evaluated for inflammation and cell death and summed for a total pathology score. Animals infected with the Spain 2009, Senegal 2003, Uganda 2010, and South Africa 1959 strains of Usutu virus generally had higher amounts of inflammation and necrosis in the liver than those infected with the Netherlands 2016 strain (Fig. 4B). Inflammation when present was typically a mix of macrophages and neutrophils (Fig. 4E). Livers taken from PBS control animals were characterized by no evidence of inflammation or cell death. Prominent cell death (arrow) and variable inflammation was identified in the liver tissues of animals infected with the Uganda 2010 isolate. Mild inflammation was identified in the liver tissues of animals infected with the Netherlands 2016 isolate (Fig. 4E). No significant microscopic lesions were identified in the heart or brain of any animal (Fig. 4F).

Due to the high viremia on the day of euthanasia, to corroborate viral titers, USUV antigen was detected in tissues by immunohistochemistry. In the spleen, animals infected with the Spain 2009, Senegal 2003, Uganda 2010, and South Africa 1959 strains of Usutu virus had large amounts of antigen within macrophages while those infected with the Netherlands 2016 strain had very minimal antigen within rare macrophages (Fig. 4D). No staining was observed in tissues from PBS control mice. Staining was revealed in macrophages within the spleen of the mice inoculated with the Uganda 2010 isolate. Rare antigen was present in macrophages in spleens from those animals infected with the Netherlands 2016 isolate. In the liver, immunohistochemistry showed variable amounts of antigen across the strains were present in Kupffer cells and inflammatory macrophages. Immunohistochemistry of the heart and brain revealed variable staining within mononuclear cells and cardiomyocytes of the heart. Thus, the immunohistochemistry supports the viral titers in the spleen.

### European USUV strains with differential pathogenesis are derived from separate introductions into Europe

To identify genetic differences that may be responsible for the differences in pathogenesis generated by USUV isolates in mice, we sequenced the five USUV isolates. The complete list of amino acid differences is available in Supplementary Table 1.

A maximum likelihood tree was constructed to evaluate the phylogenetic relationship of these isolates (Figure 5). The five USUV isolates used in this study are highlighted in blue and marked with an orange star. From this analysis, it is apparent that the Spain 2009 and Netherlands 2016 strains are derived from separate introductory events of USUV into Europe. Eighteen non-synonymous differences are unique to the Netherlands 2016 isolate compared to the other 4 isolates sequenced and tested in this study (Table 2). One or more of these distinct differences may be critical for attenuation of Netherlands 2016 in an *Ifnar*^*-/-*^ mouse model.

**Table 2.**
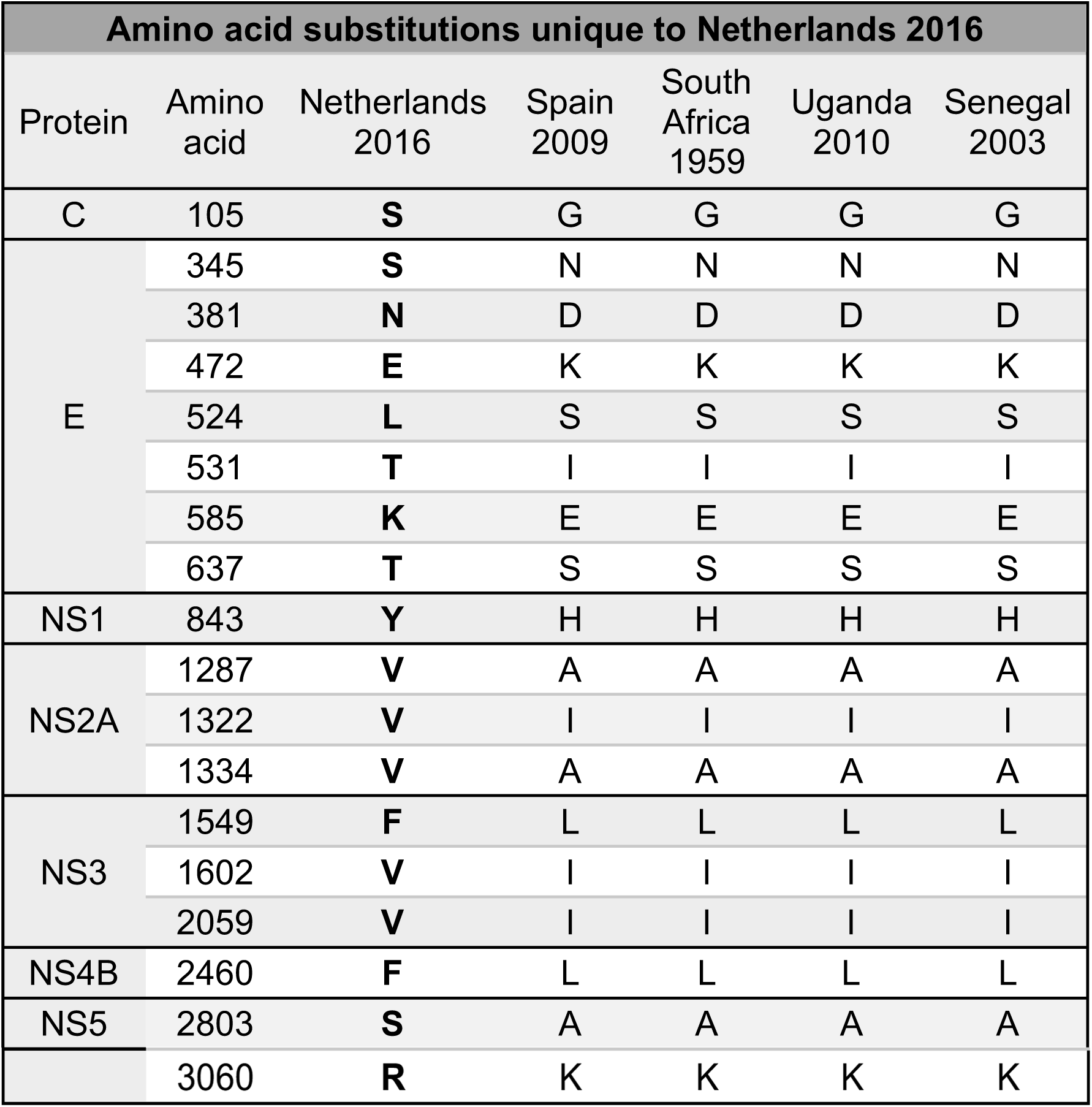

**Figure 5.**
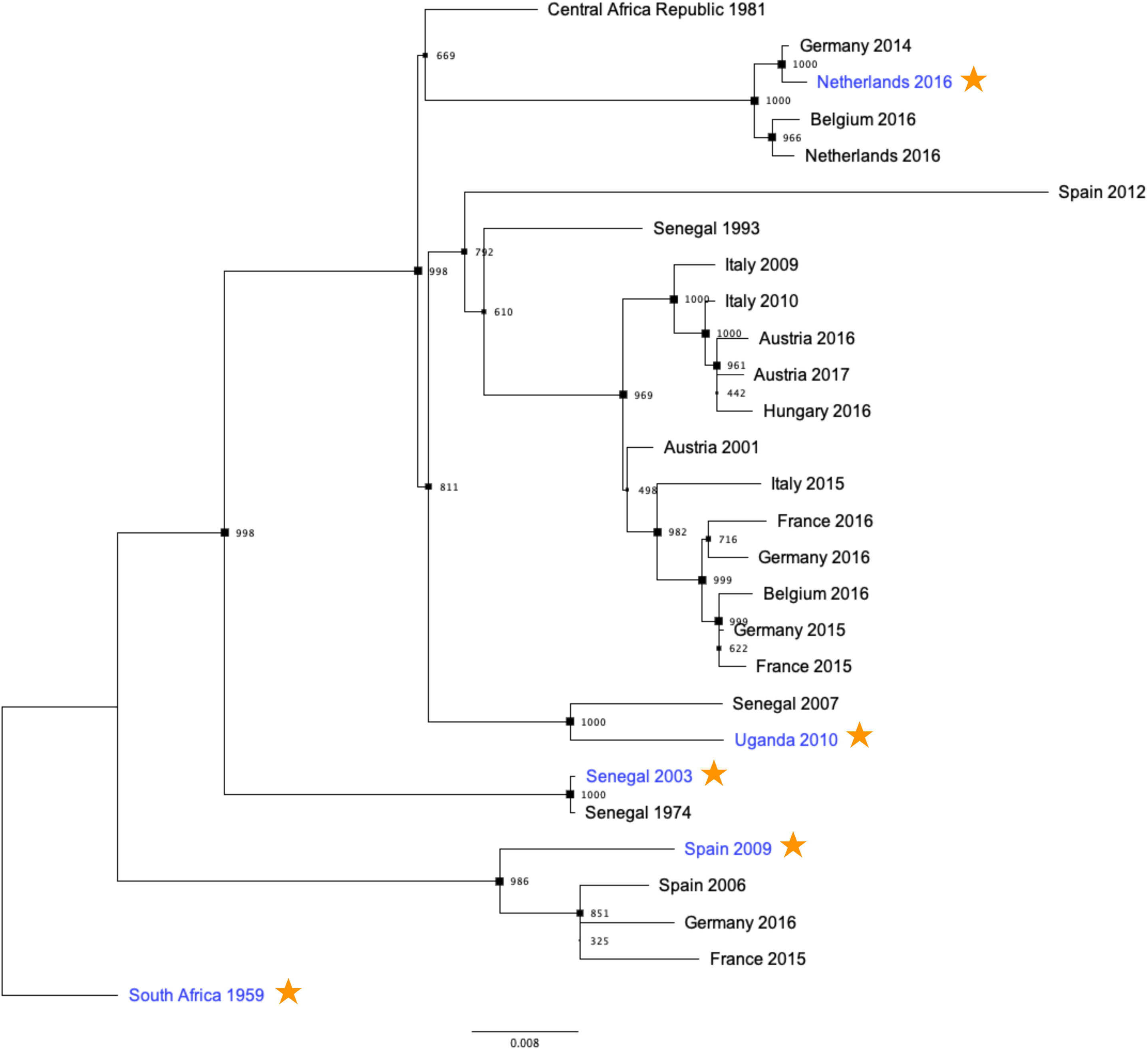
Phylogenetic analysis of African and European USUV strains. A maximum likelihood phylogenetic tree was generated using the USUV strains tested in experimental inoculations (denoted in blue with an orange star) as well as a subset of publicly available USUV sequences. Bootstrap values correspond with node size.

## Discussion

In this study, differential pathogenesis was observed in mice inoculated with two European and three African USUV isolates. All mice infected with Uganda 2010, Senegal 2003, Spain 2009, and South Africa 1959 succumbed to disease within 6 days of inoculation (Figure 2A). In contrast, 88% of mice inoculated with Netherlands 2016 survived infection. Mice inoculated with Uganda 2010 or Senegal 2003 developed viremia that was 10,000-fold higher than mice infected with Netherlands 2016 and 100-fold higher than mice infected with Spain 2009 or South Africa 1959 (Figure 3A). Spleen tissues from mice inoculated with the Netherlands 2016 strain had significantly lower levels of inflammation and cell death than other strains. These differences are likely due to several non-synonymous differences between the USUV isolates (Table 2).

The phylogenetic tree of USUV (Figure 4) indicates that USUV has been introduced into Europe at least three times, a result consistent with previous phylogenetic analyses [11, 30]. The Spain 2009 and Netherlands 2016 strains tested in this study were derived from separate introductions of USUV into Europe. Furthermore, it is apparent that USUV evolution has led to a changes in virulence, resulting in a highly pathogenic strain, such as Uganda 2010, and a less pathogenic strain, such as Netherlands 2016. Differential pathogenesis can be attributed to multiple amino acids differences across the viral genome, though even single mutations can have drastic effects on flavivirus pathogenesis. For example, a single amino acid substitution in the NS3 helicase protein in the WNV NY99 strain was found to be a significant virulence factor in American crows (*Corvus brachyrhynchos*) by performing site-directed mutagenesis [31]. Additionally, two amino acid mutations in the envelope glycoprotein of a dengue type 4 virus (DEN4) were found to be critical for neurovirulence in mice by generating intratypic chimeric viruses [32]. In order to elucidate the role of single amino acid changes in pathogenesis of USUV, future experiments could use reverse genetics to understand the effects of individual mutations on pathogenesis.

Our study showed that Netherlands 2016 is less pathogenic than the other isolates tested. This warrants further investigation into genetic determinants of attenuation in a mouse model. Netherlands 2016 and Uganda 2010, respectively, differ at residue 147 in the envelope protein and residue 157 in NS4B, both in proximity to known virulence factors of flaviviruses. Attenuation of neurovirulent WNV in mice has been associated with loss of glycosylation at residue 154 of the envelope protein [33] and a single amino acid substation at residue 102 in NS4B protein [34]. Mutation E138K in the envelope protein of Japanese encephalitis virus have also been well characterized with neurovirulence attenuation in mouse models [35, 36]. Alternatively, the attenuation of Netherlands 2016 may be due to changes occurring in the virus stock during the one passage in Vero cells that we performed in the lab to generate virus stocks or the host source of this virus strain, which was a Eurasian blackbird. Or, this phenotype may be dependent on the mouse strain used, as other flaviviruses such as Zika virus show differences in virulence between mouse strains.

Previous studies addressing USUV pathogenesis in immunodeficient murine models have described similar levels of pathogenesis for the South Africa 1959 strain or a recent European strain, Germany 2011 [28, 29]. It is clear that mice deficient in the type I interferon response are susceptible to USUV, thus suggesting that this immune response is necessary for host antiviral defenses. While these and our studies detected USUV RNA or infectious virus levels in the brain, our study is the first to assess histopathology and viral antigen in the brain of adult mice. Surprisingly, microscopic lesions and antigen staining were not identified in brain tissues from mice inoculated with USUV, suggesting that the increased pathogenesis of the recent African isolates is not due to increased neuroinvasion. However, this is consistent with a previous study that found low neuroinvasion of USUV in wild-type mice older than 1 week [27].

Studying the host responses that result in lower pathogenesis of the Netherlands 2016 strain in mice will be critical to the identification of additional mechanisms of reducing risk or treating human USUV infections. Studies investigating USUV vaccines or therapeutics are limited to one study evaluating the efficacy of a USUV vaccine in *Ifnar1*^*-/-*^ mice [29] and two studies investigating the use of antivirals in USUV infection [28, 40], suggesting a need for additional research into these areas. Based on our results, the variability in pathogenesis generated by different isolates in mice suggests that some strains, such as Uganda 2010, will be better candidates for assessing efficacy of therapeutics.

We have shown that four contemporary USUV strains differ in virulence and in genetics, with significant differences between strains circulating in Europe. Yet the findings reported here also indicate that, while outbreaks of USUV in humans in Europe have increased over the last two decades, the USUV strains circulating in Africa are also a major concern for human disease risk. Given our results, it is unlikely that the few USUV disease cases reported in Africa is due to lower pathogenesis of USUV circulating in Africa and is more likely due to limitations in disease surveillance and reporting in Africa. Thus, enhanced surveillance for USUV in Europe and Africa are necessary to more accurately estimate the burden of USUV disease.

## Acknowledgements

Funding for this project was provided by the Virginia-Maryland College of Veterinary Medicine Internal Research Competition grant. USUV isolate TMNetherlands was received from the European Virus Archive goes Global (EVAg) project that has received funding from the European Union’s Horizon 2020 research and innovation programme under grant agreement No. 653316. We thank the World Reference Center for Emerging Viruses and Arboviruses in Galveston, TX for USUV isolates HU10279-09 and DakPM173701, and the CDC in Fort Collins, CO for USUV isolates UG09615 and SAAR1776. We thank VT Laboratory Animal Research staff for contributions to animal husbandry.

## Materials and Methods

### Viruses

The USUV isolates used throughout the study were: HU10279-09 (Spain 2009) [41], TMNetherlands (Netherlands 2016) [42], UG09615 (Uganda 2010) [43], DakPM173701 (Senegal 2003) [22], and SAAR1776 (South Africa 1959) [10]. The panel of strains was chosen to assess pathogenesis of recent USUV isolates in comparison with the prototype strain, South Africa 1959. These five strains were chosen based on year of collection and availability in the United States. Virus stocks were passaged once in Vero cells upon receipt in our lab. For each isolate, RNA was extracted using the QIAamp Viral RNA Mini Kit (Qiagen). Seven overlapping RT-PCR fragments were generated with the One-Step RT-PCR Kit (Qiagen) and purified using the Zymoclean Large Fragment DNA Recovery kit (Zymo Research). Fragments were sent for inhouse Sanger sequencing (Genomics Sequencing Center, Fralin Life Sciences Institute at Virginia Tech). Primer sequences are available upon request. Sequences are available from Genbank (Accession numbers: MN813488-MN813492).

### Phylogenetic analyses

Twenty-three previously sequenced USUV genomes were selected from GenBank, with isolation dates ranging from 1981 – 2016. The coding regions of these USUV genomes and the five isolates from this study were aligned in ClustalOmega [44]. A maximum likelihood phylogenetic tree was generated in PhyML 3.0 [45] with 1000 bootstraps, and visualized using FigTree v1.4.4.

### Cell lines and growth curves

Vero cells were maintained at 37°C with 5% CO2 and cultivated in DMEM with 5% FBS and 1% penicillin-streptomycin. Vero cells were plated in 12-well plates at a density of 1.5×10^5^ cells per well and inoculated one day later at an MOI of 0.1, in triplicate. Serial time points were collected every 24 hours for 4 days. This was repeated twice for a total of 3 replicates. Samples were titrated by plaque assay in Vero cells.

### Inoculation of mice

Interferon α/β receptor 1 knockout (*Ifnar1*^*-/-*^) mice, on a C57BL/6 background, were originally received from Jackson Labs and bred on site. In two biologically independent experiments, groups of 8 (4 female and 4 male) 8-week-old mice were subcutaneously inoculated via rear footpad injection with 10^3^ PFU of USUV isolates. Blood samples were collected in serum separator tubes via submandibular vein bleeds on days 1, 3, 5 and 7 post-inoculation. Mice were weighed daily and observed for clinical signs of illness, including lethargy, tremors, and weight loss. When mice exhibited ≥15% weight loss (or at 28 days post inoculation), they were euthanized by deep isoflurane anesthesia followed by cervical dislocation. Brain, heart, liver, and spleen tissues were collected at the time of euthanasia. In a separate experiment, an additional 4 mice were inoculated with TM Netherlands and euthanized on dpi 6, in order to collect tissues for comparison to other strains. Tissues were weighed and suspended in equal parts BA-1 medium, then homogenized via bead homogenization in a Qiagen TissueLyserLT at 50 oscillations/sec for 2 minutes followed by centrifugation to pellet solids. All samples were titrated by Vero cell plaque assay.

### Histopathology and immunohistochemistry

Following euthanasia, tissues were collected and stored in 10% buffered formalin prior to standard processing and paraffin embedding. Sections were cut at 5 μm and stained by routine hematoxylin-eosin (H&E) for histopathological analysis or by immunohistochemistry (IHC) as follows. Following deparaffinization, heat induced antigen retrieval was accomplished by immersing slides in 10 mM sodium citrate buffer (pH 6.0) for 15 minutes. Slides were stained using the Pierce Peroxidase Detection Kit (Thermo Scientific). The primary antibody was the pan-flavivirus clone D1-4G2-4-15. PBS-inoculated mice served as negative controls for IHC and H&E. For immunostaining, negative control slides were prepared using tissues from PBS-inoculated mice as described above, as well as tissues from infected mice without primary antibody. Slides were analyzed by a board-certified pathologist.

### Statistical analyses

Survival curves were analyzed by Mantel-Cox test and s erum titers were analyzed by mixed-effects analysis with Tukey’s multiple comparisons test. Tissue titers were compared by ordinary one-way ANOVA with Tukey’s multiple comparisons test. Growth curves were analyzed with a two-way ANOVA with Tukey’s multiple comparisons test. All statistical analyses were performed in GraphPad Prism 8.

### Ethics Statement

All studies were conducted under approved IACUC protocol #18-085 at Virginia Tech.

**Supplementary Table 1.**

| **Amino acid substitutions compared to South Africa 1959** |  |  |  |  |  |  |
| --- | --- | --- | --- | --- | --- | --- |
| Protein | Amino acid | South Africa 1959 | Spain 2009 | Netherlands 2016 | Uganda 2010 | Senegal 2003 |
| **C** | 101 | K | **R** | K | K | K |
|  | 105 | G | G | **S** | G | G |
|  | 112 | V | **L** | **L** | V | V |
|  | 123 | M | **I** | M | M | M |
| **prM** | 153 | T | **M** | T | T | T |
|  | 172 | D | **G** | D | D | D |
|  | 215 | R | **L** | R | R | R |
|  | 283 | V | V | V | V | **I** |
| **E** | 345 | N | N | **S** | N | N |
|  | 381 | D | D | **N** | D | D |
|  | 386 | K | **R** | K | K | K |
|  | 472 | K | K | **E** | K | K |
|  | 506 | V | V | V | **I** | V |
|  | 524 | S | S | **L** | S | S |
|  | 531 | I | I | **T** | I | I |
|  | 569 | S | **G** | **G** | **G** | **G** |
|  | 585 | E | E | **K** | E | E |
|  | 595 | G | **S** | G | G | G |
|  | 613 | S | **G** | **G** | **G** | **G** |
|  | 637 | S | S | **T** | S | S |
|  | 790 | S | **N** | **N** | **N** | **N** |
| **NS1** | 843 | H | H | **Y** | H | H |
|  | 891 | K | K | K | **R** | **R** |
|  | 900 | T | T | T | T | **I** |
|  | 939 | V | V | V | V | **A** |
|  | 967 | H | **Y** | H | H | H |
|  | 1067 | V | **I** | V | V | V |
|  | 1117 | K | **R** | **R** | **R** | **R** |
| **NS2A** | 1190 | I | I | I | I | **V** |
|  | 1227 | A | **V** | A | A | A |
|  | 1236 | A | **T** | A | A | A |
|  | 1240 | L | L | L | L | **F** |
|  | 1267 | D | **N** | **N** | **N** | **N** |
|  | 1268 | L | **F** | L | **F** | **F** |
|  | 1270 | L | **F** | L | L | L |
|  | 1287 | A | A | **V** | A | A |
|  | 1322 | I | I | **V** | I | I |
|  | 1334 | A | A | **V** | A | A |
| NS2B | 1436 | T | **A** | T | T | **A** |
|  | 1460 | I | **V** | I | I | I |
|  | 1492 | I | I | I | I | **V** |
| NS3 | 1549 | L | L | **F** | L | L |
|  | 1602 | I | I | **V** | I | I |
|  | 1618 | V | V | **I** | **I** | **I** |
|  | 1645 | K | **R** | K | K | K |
|  | 1981 | G | **S** | G | G | G |
|  | 1983 | S | S | S | **N** | S |
|  | 2059 | I | I | **V** | I | I |
|  | 2075 | I | **V** | I | I | I |
|  | 2106 | S | S | **A** | **A** | S |
| NS4B | 2290 | G | **S** | **S** | **S** | **S** |
|  | 2294 | P | **S** | P | P | P |
|  | 2301 | P | **H** | P | P | P |
|  | 2355 | N | **T** | N | N | N |
|  | 2393 | T | **I** | T | T | T |
|  | 2460 | L | L | **F** | L | L |
| NS5 | 2550 | R | R | **K** | **K** | R |
|  | 2552 | D | **E** | D | D | D |
|  | 2620 | A | **V** | A | A | A |
|  | 2695 | E | **D** | E | E | E |
|  | 2706 | R | R | R | R | **K** |
|  | 2765 | T | **P** | T | T | T |
|  | 2777 | T | T | T | T | **A** |
|  | 2784 | E | **D** | E | E | E |
|  | 2803 | A | A | **S** | A | A |
|  | 2849 | G | **S** | **S** | **S** | **S** |
|  | 2902 | R | **K** | R | R | R |
|  | 3060 | K | K | **R** | K | K |
|  | 3088 | M | **I** | M | M | M |
|  | 3322 | V | **I** | V | V | V |
|  | 3430 | E | E | E | E | **G** |

## Notes

### Competing Interest Statement

The authors have declared no competing interest.

